# Syncollin secreted by activated human neutrophils targets bacteria

**DOI:** 10.1101/2022.02.14.480106

**Authors:** Rosie A. Waters, James Robinson, J. Michael Edwardson

## Abstract

Syncollin is a 16-kDa protein that was originally isolated from the pancreatic zymogen granule. Syncollin is also found in human neutrophils and is secreted from promyelocytic HL-60 cells upon stimulation. Recently, we reported that syncollin is able to bind to bacterial peptidoglycan, to damage the bacterial envelope and to restrict bacterial growth. Here, we show that syncollin is secreted from activated primary human neutrophils, and interacts with extracellular neutrophil traps (NETs) in a similar manner to the well-characterized granular protein myeloperoxidase. In addition, secreted syncollin is able to coat the surface of *E. coli* in co-cultures of neutrophils and bacteria. On the basis of our findings, we suggest that syncollin plays a role in host defence in the blood.

## Introduction

Syncollin is a 16-kDa protein that was originally isolated from the pancreatic zymogen granule (1). Syncollin is also found in human neutrophils and in promyelocytic HL-60 cells; it is present in the azurophilic granules of the latter, and is secreted from these cells upon stimulation (2). Syncollin is able to oligomerize, and forms doughnut-shaped structures on lipid bilayers (3). Further, syncollin is able to permeabilize both erythrocytes (4) and liposomes (3). Recently, we showed that syncollin binds to bacterial peptidoglycan, and restricts the growth of both Gram-positive and Gram-negative bacteria (5). Additionally, we found that syncollin was able to permeabilize *E. coli* membranes and to cause surface damage to both *L. lactis* and *E. coli*. In the present study, we explored the behaviour of syncollin secreted from human neutrophils. We show that secreted syncollin interacts with extracellular neutrophil traps (NETs) and coats the surface of exogenously added *E. coli*. We suggest that syncollin plays a role in host defence not only in the gut, as previously described (5), but also in the blood.

## Materials and methods

### Isolation of human neutrophils

All reagents were from Merck (Gillingham, Dorset), unless otherwise indicated. Human blood (25 mL) was layered gently onto Histopaque-1077™ (15 mL) and centrifuged at 1450 rpm for 30 min. The layer above the erythrocyte-rich fraction, containing mononuclear cells, was removed and the remaining erythrocyte-rich layer was diluted with an equal volume of Hanks' balanced salt solution without Ca^2+^ or Mg^2+^. The cell suspension was further diluted with an equal volume of 2% (w/v) dextran, gently mixed by inversion and left for 30 min at room temperature to allow the erythrocytes to sediment. The erythrocyte-free top layer was mixed with an equal volume of balanced salt solution and centrifuged at 1150 rpm for 5 min. The supernatant was gently removed and the neutrophil pellet was resuspended in Roswell Park Memorial Institute (RPMI) medium containing 10% (v/v) fetal bovine serum (FBS) and penicillin/streptomycin. (If neutrophils were to be incubated with bacteria, the neutrophil pellet was resuspended in medium without antibiotics).

### Immunofluorescence imaging of neutrophils

Sterile fibronectin (10 ng.mL^−1^) was added to flat-bottomed imaging dishes (Ibidi; Thistle Scientific, Glasgow) and left to incubate for 2 h at 37°C. Wells were washed twice with sterile phosphate-buffered saline (PBS), pH 7.0, and dried in air. Neutrophils (~2 × 10^6^ cells.mL^−1^ in a 30-μL volume) were added to the plates and left to adhere at 37°C for 30 min under sterile conditions. Where appropriate, neutrophils were activated by addition of 200 nM *N*-formylmethionyl-leucyl-phenylalanine (fMLP) in RPMI medium containing 10% (v/v) fetal bovine serum (FBS), penicillin/streptomycin for 30 min at 37°C. Quiescent or activated cells were fixed by incubation in sterile 4% (w/v) paraformaldehyde for 20 min at room temperature and then washed three times with 30 μL PBS. To permeabilize cells, 30 μL of 0.1% (v/v) Triton X-100 was added to each well, followed by incubation for 5 min and three gentle washes with PBS. Neutrophil Fc receptors were blocked by incubation with 10 μg.mL^−1^ Fc-Receptor block (BD, Wokingham, Berkshire) in PBS containing 1% (w/v) bovine serum albumin (BSA) for 1 h at room temperature. Primary antibody (rabbit polyclonal anti-syncollin (6) or mouse monoclonal anti-myeloperoxidase [MPO; Santa Cruz Biotechnology, Heidelberg, Germany]) diluted in PBS containing 1% (w/v) BSA was then added to each well for overnight incubation at 4°C. Cells were washed gently three times with PBS and secondary antibody (fluorescein isothiocyanate-[FITC]-conjugated anti-rabbit or allophycocyanin-[APC]-conjugated anti-mouse), again in PBS containing 1% (w/v) BSA, was added to each well and left for 2 h at room temperature in the dark. Hoechst 33342 (Thermo Fisher, Loughborough, Leicestershire; 1 μg.mL^−1^ in PBS) was also added for detection of DNA. Control samples, exposed to secondary antibody alone, were also included. After a final three washes with PBS, immersion oil (Ibidi) was added to each well and samples were imaged using a Leica (Milton Keynes, Buckinghamshire) SP5 confocal microscope with constant exposure time, brightness and contrast adjustments for each experiment. Cells in immunofluorescence images were defined and fluorescence intensity in each cell area was quantified (in arbitrary units [A.U.]) using ImageJ Fiji™. Two cell wells were analyzed for each condition in order to generate the intensity datapoints.

### Analysis of neutrophil activation by immunoblotting

A neutrophil suspension (5 × 10^6^ cells.mL^−1^) was incubated for 30 min at 37°C either with or without 200 nM fMLP. Neutrophils were then centrifuged at 5000 rpm for 3 min. Pellets and supernatants (after methanol-chloroform precipitation) were analysed by sodium dodecyl sulphate-polyacrylamide gel electrophoresis and immunoblotting using a rabbit polyclonal anti-syncollin antibody (Hodel and Edwardson, 2000) followed by a horseradish peroxidase-conjugated goat anti-rabbit antibody and enhanced chemiluminescence imaging (Thermo Fisher). Band intensities were quantified (in A.U.) using ImageJ™.

### Interaction of recombinant syncollin with *E. coli*

Syncollin bearing a double-Strep-II tag at its C-terminus was isolated from the supernatant of tsA-201 cells through binding to Strep-Tactin XT™ beads (IBA, Göttingen, Germany), as described previously (5). Strep-tagged syncollin was incubated with *E. coli* for 3 h at 4°C. Bacteria were then pelleted by centrifugation and washed three times with PBS. Bacteria were added to imaging dishes (Ibidi) and incubated with a rabbit polyclonal anti-syncollin antibody for 2 h at room temperature followed by an FITC-conjugated goat anti-rabbit secondary antibody and 4′,6-diamidino-2-phenylindole (DAPI, Vector Laboratories, Burlingame, CA; 1.5 μg.mL^−1^).

### Fluorescence imaging of neutrophils after activation by bacterial infection

An *E. coli* suspension at an optical density (660 nm) of 1.5 was added in 30-μL volumes to poly-L-lysine-coated flat-bottomed imaging dishes (Ibidi™) and allowed to adhere for 30 min at 37°C. Medium was then removed before addition of 30 μL of freshly isolated neutrophils at 1.6 × 10^6^ cells.mL^−1^ followed by incubation at 37°C for 60 and 120 min. At each time point, medium was removed from a well and the cells were fixed and permeabilized as described above. Cells were washed twice with PBS before addition of blocking buffer (PBS containing 10% [v/v] FBS, 3% [w/v] BSA and Fc-Receptor blocker) followed by overnight incubation at 4°C. Cells were then incubated overnight at 4°C with primary antibodies (rabbit polyclonal anti-syncollin and mouse monoclonal anti-myeloperoxidase [MPO]). Wells were washed three times with PBS before addition of secondary antibodies (fluorescein isothiocyanate- [FITC]-conjugated anti-rabbit and allophycocyanin- [APC]-conjugated anti-mouse), and DAPI (1.5 μg.mL^−1^) followed by incubation for 2 h at room temperature in the dark. Samples were imaged using confocal microscopy.

## Results and Discussion

### Syncollin is secreted from activated neutrophils

Immunofluorescence detection of syncollin in resting neutrophils revealed a punctate distribution throughout the cytoplasm (Fig. 1A). Activation of the neutrophils by treatment with 200 nM fMLP for 30 min caused them to become less uniformly round and more flattened; syncollin also became more localized near the plasma membrane. Syncollin fluorescence intensity in individual cells was measured before and after activation. As a control for secretion of granular proteins, the staining of MPO was also quantified. As shown in Fig. 1B, and quantified in Fig 1C, the fluorescence intensity of syncollin and MPO fell in a similar way after a 30-min activation, consistent with the secretion of both proteins.

**Fig. 1.**
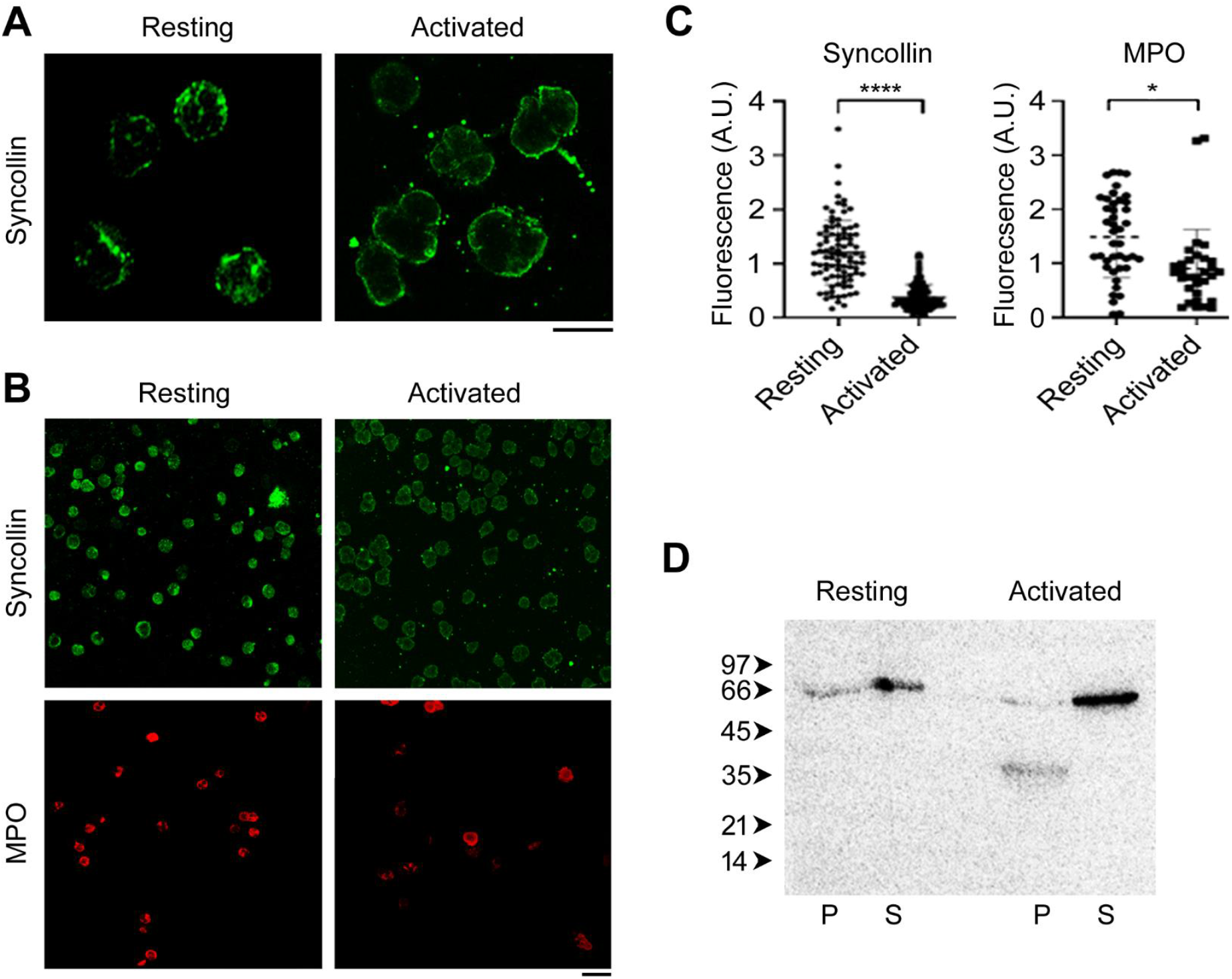
Syncollin is secreted from activated neutrophils. (A) Immunofluorescence detection of syncollin in resting and activated neutrophils. Neutrophils were activated by incubation with 200 nM fMLP for 30 min at 37°C and then fixed and permeabilized. Syncollin was detected using a rabbit polyclonal primary antibody followed by an FITC-conjugated goat anti-rabbit secondary antibody. Scale bar, 10 μm. (B, C) Quantitation of syncollin and MPO secretion from activated neutrophils. Cells in immunofluorescence images (B) were defined and fluorescence intensity (in A.U.) was quantified using ImageJ Fiji™ (C). The numbers of cells analysed were: syncollin resting, 92; syncollin activated, 109; MPO resting, 42; MPO activated, 31. Scale bar, 10 μm. ****, p<0.0001; *, p<0.05 (Student’s unpaired t-test). (D) Immunoblot of cell pellet (P) and supernatant (S) fractions of resting and activated neutrophils. Molecular mass markers (kDa) are shown at the left. Band intensities were quantified using ImageJ™, and are (in A.U.): resting pellet, 1616; resting supernatant, 3785; activated pellet, 621+1278=1899; activated supernatant, 9023.

Pellets and supernatants were collected from resting and activated neutrophils and immunoblotted for syncollin. In resting cells, syncollin migrated as a ~65 kDa band, indicating the formation of very stable oligomers (likely tetramers), as observed previously (2, 6) Interestingly, the major band in activated cells ran at ~35 kDa, indicating that the oligomer may become less stable upon activation. Quantitation of the bands revealed that, consistent with the immunofluorescence result, the syncollin level in the neutrophil pellet fell from 30% of total in resting neutrophils to 17% of total in activated neutrophils, with a corresponding increase in the supernatant from 70% to 83% of total (Fig. 1D).

### Secreted syncollin binds to NETs

Neutrophils were next activated by addition of exogenous bacteria (*E. coli*) in an attempt to generate an *in-vitro* model of a bacterial infection. Neutrophils were allowed to adhere to fibronectin-coated imaging plates and a suspension of *E. coli* was then added. The neutrophils were then fixed, permeabilized and subjected to immunofluorescence analysis. Neutrophil extracellular traps (NETs) began to appear after incubation of the neutrophils with *E. coli* (*arrows* in Fig. 2). The NETs could be stained with Hoechst 33342, as expected, given that they contain de-condensed neutrophil DNA (7). As shown, both syncollin and MPO were also present in the NETs.

**Fig. 2.**
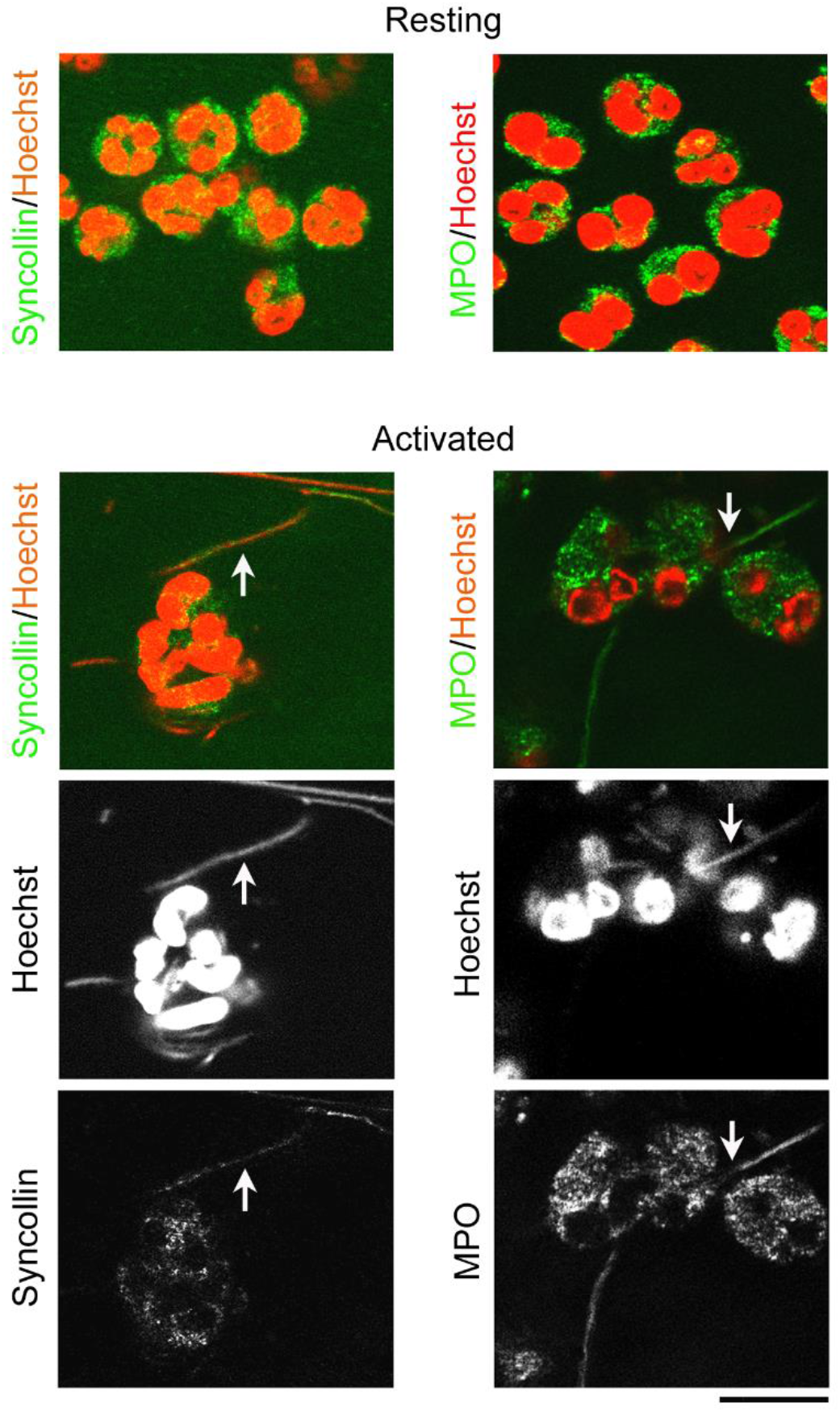
Secreted syncollin binds to NETs. Neutrophils were allowed to adhere to a fibronectin-coated surface and either left in a resting state or activated by addition of a suspension of *E. coli* followed by incubation at 37°C for 45 min. Cells were then fixed and permeabilized. Syncollin was detected using a rabbit polyclonal primary antibody followed by an FITC-conjugated secondary antibody (green). MPO was detected using a monoclonal primary antibody followed by an APC-conjugated secondary antibody (green). DNA was detected using Hoechst 33342 (red). Individual channels are also presented in black and white to reveal the co-localization of both syncollin and MPO with NET DNA. *Arrows* indicate NETs. Scale bar, 20 μm.

### Recombinant Strep-tagged syncollin binds to bacteria

The inhibitory actions of recombinant syncollin on bacterial growth have been reported recently (5), and it was suggested that syncollin must bind to the bacterial capsule to mediate its inhibitory action. To test this idea, bacteria (*E. coli*) were incubated with recombinant Strep-tagged syncollin for 3 h at 4°C, washed three times and then subjected to immunofluorescence imaging with anti-syncollin antibody. DAPI staining revealed the presence of the bacteria (Fig. 3), and syncollin could be seen bound to the bacterial surface consistent with its ability to bind to peptidoglycan and to restrict bacterial growth (5). A faint background fluorescence signal was seen when FITC-conjugated secondary antibody was used alone.

**Fig. 3.**
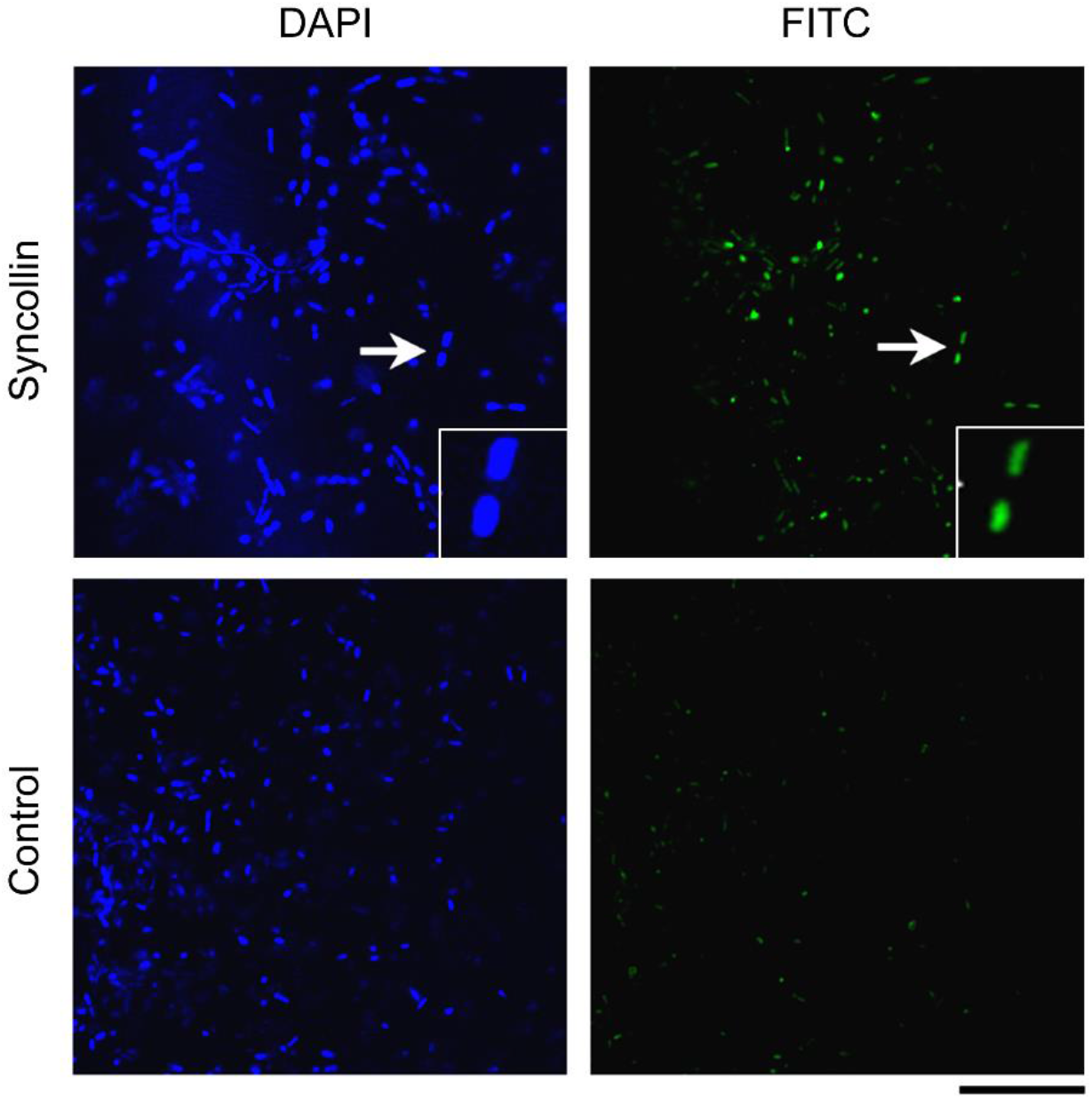
Recombinant Strep-tagged syncollin binds to bacteria. Purified syncollin-Strep (0.05 mg/mL) or buffer alone (control) were incubated with *E. coli* for 3 h at 4°C. Bacteria were washed and incubated with a polyclonal anti-syncollin primary antibody followed by FITC-conjugated secondary antibody (green), and DAPI, to stain DNA (blue). Scale bar, 20 μm. Insets show zoomed images of syncollin-decorated bacteria, indicated by *arrows* in the main panels.

### Syncollin secreted by activated neutrophils decorates exogenously-added bacteria

Having shown that *E. coli* can be decorated by recombinant Strep-tagged syncollin, we next asked whether endogenous syncollin released from activated neutrophils would also bind to the bacteria. *E. coli* were allowed to adhere to fibronectin-coated imaging dishes, and neutrophils (20 μL at 3.6 × 10^6^ cells/mL) were added. After 60 and 120 min, cells were fixed, permeabilized and subjected to immunofluorescence imaging with anti-syncollin and anti-MPO antibodies. Consistent with the results described above, syncollin could clearly be seen in association with the bacteria at both time-points, although the staining appeared more obvious after 120 min (*arrows* in Fig. 4). In contrast, MPO did not decorate the bacteria.

**Fig. 4.**
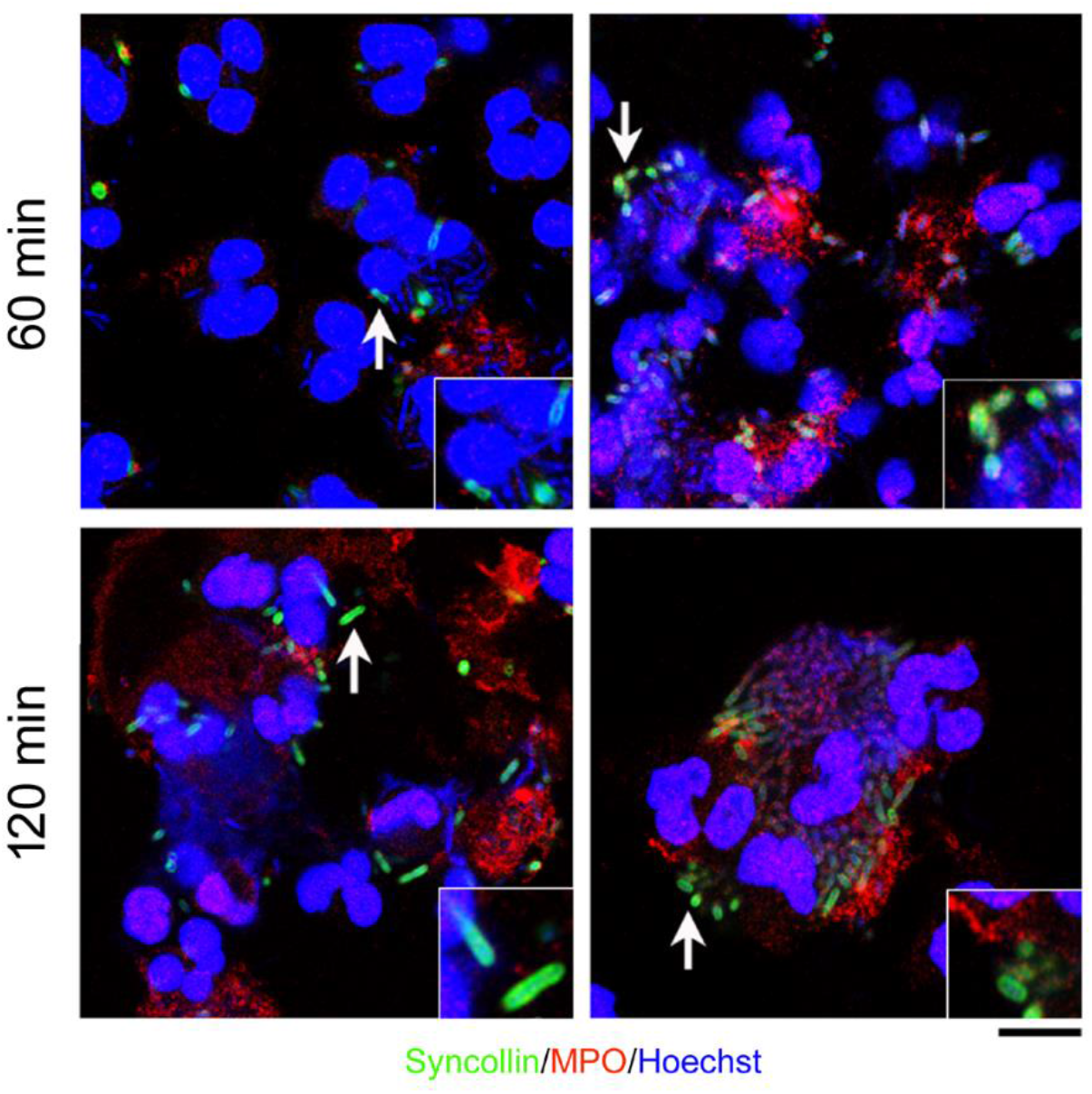
Syncollin secreted by activated neutrophils decorates exogenously-added bacteria. Neutrophils were added to adherent *E. coli* and immunofluorescence images were taken after 60 min (*top*) and 120 min (*bottom*) of activation. Syncollin was detected using a polyclonal primary antibody followed by an FITC-conjugated secondary antibody (green). MPO was detected using a monoclonal primary antibody followed by an APC-conjugated secondary antibody (red). Hoechst 33342 was added to detect DNA (blue). Syncollin-decorated bacteria are indicated by *arrows*. Scale bar, 10 μm. Insets show zoomed images of syncollin-decorated bacteria, indicated by *arrows* in the main panels.

It has been reported previously (2) that syncollin is expressed in neutrophils and HL-60 cells, likely in the azurophilic granules, and is released from the latter upon activation, hinting at a role in host defence. Here, we show that syncollin is secreted from the neutrophils upon activation. The secreted syncollin decorates NETs, and also binds to the surface of exogenously added bacteria, as demonstrated through the use of a co-culture model. NETs are weblike structures consisting of decondensed chromatin to which a number (>20) of cytosolic and granular proteins bind (7). NETs trap and kill invading microorganisms, including bacteria, fungi and viruses. MPO binding to NETs has been demonstrated previously (8); however, the presence of syncollin in these structures is a novel finding. We suggest that in a physiological setting, syncollin is released from the neutrophils in response to a bacterial challenge, and then binds to bacteria to produce an anti-microbial effect.

## Abbreviations

APC: allophycocyanin
A.U.: arbitrary units
BSA: bovine serum albumin
DAPI: 4′,6-diamidino-2-phenylindole
FBS: fetal bovine serum
FITC: fluorescein isothiocyanate
fMLP: N-formylmethionyl-leucyl-phenylalanine
MPO: myeloperoxidase
NET: neutrophil extracellular trap
PBS: phosphate-buffered saline
RPMI: Roswell Park Memorial Institute.

## Acknowledgements

We are grateful to Rebecca Riddle, Sarah Millington-Burgess and all other members of the Matthew Harper laboratory (Department of Pharmacology, University of Cambridge) for providing neutrophils and advice about their use. RAW was supported by an AstraZeneca Graduate Studentship and a David James Studentship from the Department of Pharmacology.

## Author contributions

RAW, JR and JME designed the study. RAW conducted the experiments and analysed the data. All three authors wrote the manuscript.

